# OpenSense: An open-source toolbox for Inertial-Measurement-Unit-based measurement of lower extremity kinematics over long durations

**DOI:** 10.1101/2021.07.01.450788

**Authors:** Mazen Al Borno, Johanna O’Day, Vanessa Ibarra, James Dunne, Ajay Seth, Ayman Habib, Carmichael Ong, Jennifer Hicks, Scott Uhlrich, Scott Delp

## Abstract

**Background:** The ability to measure joint kinematics in natural environments over long durations using inertial measurement units (IMUs) could enable at-home monitoring and personalized treatment of neurological and musculoskeletal disorders. However, drift, or the accumulation of error over time, inhibits the accurate measurement of movement over long durations. We sought to develop an open-source workflow to estimate lower extremity joint kinematics from IMU data that was accurate, and capable of assessing and mitigating drift.

**Methods:** We computed IMU-based estimates of kinematics using sensor fusion and an inverse kinematics approach with a constrained biomechanical model. We measured kinematics for 11 subjects as they performed two 10-minute trials: walking and a repeated sequence of varied lower-extremity movements. To validate the approach, we compared the joint angles computed with IMU orientations to the joint angles computed from optical motion capture using root mean square (RMS) difference and Pearson correlations, and estimated drift using a linear regression on each subject’s RMS differences over time.

**Results:** IMU-based kinematic estimates agreed with optical motion capture; median RMS differences over all subjects and all minutes were between 3-6 degrees for all joint angles except hip rotation and correlation coefficients were moderate to strong (r = 0.60 to 0.87). We observed minimal drift in the RMS differences over ten minutes; the average slopes of the linear fits to these data were near zero (−0.14 to 0.17 deg/min).

**Conclusions:** Our workflow produced joint kinematics consistent with those estimated by optical motion capture, and could mitigate kinematic drift even in the trials of continuous walking without rest, obviating the need for explicit sensor recalibration (e.g. sitting or standing still for a few seconds or zero-velocity updates) used in current drift-mitigation approaches. This could enable long-duration measurements, bringing the field one step closer to estimating kinematics in natural environments.

## Introduction

Inertial measurement units (IMUs) could enable biomechanics and rehabilitation researchers to measure kinematics in a variety of populations, in natural environments and over long durations. From detecting functional improvement in patients post-stroke to monitoring fall-risk in older adults (1), continuous sensing of kinematics could improve our understanding of human movement pathology by providing many repetitions of a movement in home or community settings, in contrast with the limited number of trials and highly-controlled environment of a laboratory experiment. IMUs could also enable the early detection of disease or injury-risk. They could then be used together with mobile interventions to create rehabilitation or injury-prevention strategies that are optimized to the user’s biomechanics. In addition, IMUs are an inexpensive way to measure movement in large cohorts, facilitating large-scale multicenter clinical trials for which traditional motion capture is currently infeasible.

IMUs have been used to estimate kinematics during human movement for the past 30 years (2) and over the past decade, the biomechanics and rehabilitation communities have substantially improved the accuracy of IMU-based methods for measuring kinematics. For example, researchers have developed new sensor fusion algorithms to estimate orientations (3–6) and devised more precise sensor-to-body segment alignment methods (7,8). Researchers have also employed biomechanical models (9–11), and used neural networks and optimization (12–15) to estimate accurate kinematics without reliance on potentially distorted magnetometer data or precise IMU placement. While studies have shown accuracy for lower extremity kinematics on the order of one degree root mean square (RMS) difference compared to optical motion capture, the overwhelming majority of studies assess accuracy of steady-state behavior (e.g., walking or running) over short durations (on the order of one minute or less) (16), even though these conditions are not wholly representative of natural behavior.

A key challenge estimating 3D orientation from IMUs over long durations is compounding drift over time. Most IMU-based algorithms to estimate joint kinematics rely on three-dimensional orientations computed through sensor fusion, a process where triaxial data from the accelerometer, gyroscope, and/or magnetometer are combined to give a more accurate measure of orientation than could be provided by any of the single data streams. Strap-down integration, or integrating gyroscope data from an IMU that is strapped to the body segment of interest (as opposed to mounted on a stabilized platform), results in random drift as numerical integration amplifies noise in the gyroscope data (17). Accelerometer-based and magnetometer-based compensation can correct this drift using the earth’s gravitational and magnetic field vector. However, ferromagnetic disturbances distort the measurement of earth’s magnetic field, which can lead to inaccurate orientation estimates (18). Sensor fusion approaches have been designed to mitigate drift in specific and precise movements such as those performed by robots (6). Validation studies applying sensor fusion methods in human movement analysis have reported average RMS differences in the range of 1.7 to 8 degrees for joint angles over short durations (on the order of one minute) (5,19–21). Recently, validation of sensor fusion approaches in human movement analysis over longer durations has emerged, though validation has focused on single joints (22,23) and assumed that the user took regular rests in the same posture each time (22). It remains undetermined whether these methods translate to other joints and activities.

The physiological joint constraints imposed by biomechanical models mitigate errors due to drift. Previous studies have computed IMU-based estimates of kinematics using biomechanical models with accuracies under 5 degrees RMS difference. For the most part, however, these studies have used closed-source commercially available models (e.g., MVN Xsens) (24) that cost on the order of $10k, or simple models that are developed in-house and thus are limited to those with IMU and modeling expertise (10,25–29). Tagliapietra et al. (2018) provides an open-source IMU-based inverse kinematics algorithm using a biomechanical model; this study reports good agreement (RMS differences less than 6 degrees) between their IMU-based estimates of kinematics and the robotic-encoder-based or optical-based kinematics, but the approach has not been tested for human movement.

Ideally, the research community would have an open-source platform to compute kinematics from experimentally recorded IMU data using a physiologically representative biomechanical model that had been evaluated for use over long durations. This integrated environment would empower researchers to generate further analyses and insights (e.g. estimations of musculotendon lengths or velocities required to generate motion) that would otherwise involve invasive and complex experiments.

Our goal was thus to develop an open-source workflow for computing three-dimensional joint kinematics with IMU sensors using a human biomechanical model, and evaluate it against optical motion capture over 10-minute periods of common activities. We also sought to investigate whether an approach using sensor fusion algorithms designed and validated for arbitrary movements with robots would be robust to drift when used to compute IMU-based estimates of kinematics during a 10-minute period of human movement. Finally, we sought to provide an open dataset of synchronized IMU and optical motion capture data.

## Methods

### Data collection

We collected IMU and optical motion capture data for 11 subjects in a laboratory environment, which included significant amounts of electronic equipment and ferromagnetic materials. All subjects provided informed consent to a protocol approved by the Stanford University Institutional Review Board. Subjects were young (26.7 ± 2.7 years, mean ± 1 standard deviation (sd) and free of any musculoskeletal injuries or disorders; the mean body mass index of subjects was in the “normal” range (20.4 ± 8.6 kg/m^2^); and the majority were male (9/11). Subjects were outfitted with 8 IMUs (MTw Awinda, Xsens North America Inc., Culver City, USA), affixed to thin plexiglass plates with clusters of at least 4 retro-reflective markers, constituting a *marker plate*, and secured to the upper back (T2), lower back (L5), and the right and left thighs, shanks, and feet (Supplementary Figure S1). IMU signals were sampled at 100 Hz for nine subjects, and at 40 Hz for two subjects (due to a protocol inconsistency). IMU data were acquired via a graphical interface (MT Studio, Xsens North America, USA).

Optical motion capture data were collected simultaneously to enable comparison of the IMU-based estimates of joint kinematics to the current gold standard. In addition to the markers on the marker plate, markers were placed on the bony landmarks of the C7 vertebrae, sternoclavicular joints, acromion processes, anterior and posterior superior iliac spines, medial and lateral femoral epicondyles, medial and lateral malleoli, calcanei, and 5th metatarsal heads. Markers on the medial femoral epicondyles and malleoli and makers obscured by the marker plates were removed prior to walking trials. Marker trajectories were measured at 100 Hz using an eight-camera motion capture system (Motion Analysis Corporation, Santa Rosa, CA, USA). A standard video camera (30 frames/s) was used to record each trial and visually confirm events or event timings offline. The optical motion capture and IMU data were synchronized by maximizing the cross-correlation between the resulting joint kinematics.

### Experimental conditions

Experimental data were collected while each subject completed two conditions: (i) 10 minutes of walking and turning and (ii) 10 minutes of a repeated series of movements. Subjects started each condition with an initial calibration pose, standing with their arms by their sides, feet hip’s width apart, and facing forward for a period of five seconds. In the first condition, subjects were instructed to walk straight for 5 meters at a self-selected pace then turn 180 degrees using a self-selected strategy and to repeat this sequence for a continuous 10-minute trial. Next, subjects took the calibration pose again, and then completed multiple cycles of lower-extremity movements for 10 minutes. Each cycle consisted of sitting, standing, ascending and descending three stairs, side-stepping for five meters, walking around a 12-m oval circuit, and finally running around a 12-m oval circuit. Subjects completed the cycle 6-10 times over the 10 minutes.

### Sensor fusion

We tested three sensor fusion algorithms: a proprietary filter (embedded on-board the Xsens IMU sensor), and two open-source complementary filters (3,5). The complementary filters used the raw accelerometer, gyroscope, and magnetometer signals read from the IMU sensors. We implemented the complementary filters using the developers’ open-source code (3,5) in MATLAB R2019a (Mathworks, Inc., Natick MA, USA) with the initial orientation estimate computed from the accelerometer and magnetometer measurements when the sensors were at rest (i.e., when the subject was standing still for a few seconds). We manually tuned the filter gain (“beta parameter”) of the complementary filters (5) using data from two randomly chosen subjects (sampled at 100 Hz), and this filter gain of 0.1 was used for all subjects with data collected at 100 Hz. For the two subjects with data collected at 40 Hz, the filter gain value of 0.1 overcorrected drift resulting in poor accuracy, so we experimented with filter gains of 0.05 and 0.025 and found the latter was optimal. To evaluate the accuracy of the sensor fusion algorithm, we compared the IMU-based orientation estimates to those computed using the motion capture markers affixed to the marker plates. The orientation difference was expressed as the angle of an axis-angle representation of the relative rotation. From this angle, we computed RMS errors. As the markers and IMUs were rigidly mounted to the marker plate, we would expect minimal errors in the optical estimates of orientation, thus we report RMS errors as the main error metric and refer to them as *sensor fusion errors*.

### Inverse kinematics workflow

We used OpenSim 4.2 (30,31) (simtk.org/projects/opensim) to compute both IMU-based and optical-motion-capture-based estimates of kinematics, which we refer to as IMU-based kinematics and optical-based kinematics, respectively. We used a physiological skeletal model with 22 segments and 43 degrees-of-freedom (dofs). The model had 16 dofs in the lower body including 6 for the pelvis and 5 for each lower extremity. The hip was modeled as a ball-and-socket joint (3 dofs), the knee as a custom joint with 1 dof (32–34). The 3-dof ankle joint in the Rajagopal model was simplified to a pin joint representing ankle plantarflexion-dorsiflexion to decrease computational complexity for the inverse kinematics solver and therefore decrease computation time. The model was scaled to match each subject’s anthropometry based on experimentally-measured markers placed on anatomical landmarks. Model scaling was only relevant for computing optical-based kinematics, as rigid body segment length does not affect IMU-based inverse kinematics. For optical-based kinematics, we used an inverse kinematics algorithm to solve for the joint angles that minimized the difference between the experimentally measured marker positions and the corresponding virtual markers on the model.

Our OpenSense toolkit in OpenSim 4.2 was used to compute IMU-based joint kinematics. The IMU orientation data resulting from a given sensor fusion algorithm were imported and associated with a rigid body (e.g., pelvis) based on a user-defined sensor mapping. To determine the orientation of the IMUs relative to the body segment on which they were placed, we used the calibration pose data. We used the optical motion capture data to pose the model (given that our main focus was assessing drift) and the IMU calibration data to compute the orientations of the IMUs relative to the posed model’s body segments as fixed rotational offsets. The modeled virtual IMU frames were then assigned these offsets relative to the underlying rigid body.

After this calibration step, we used Eq. 1 to compute the difference, expressed as the angle (*θ_i_*) of the axis-angle representation between the experimentally measured IMU orientations (a rotation matrix expressed in the Earth’s reference frame *N*, 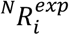), and the orientations of the model’s virtual IMUs 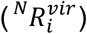. We denote this difference as *R_i_*. We used an inverse kinematics algorithm (Eq. 2) that solved for the joint angles (*q*) that minimized this weighted-squared difference 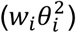.

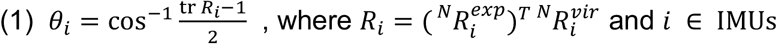

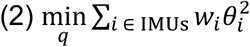

We used *θ_i_* to quantify the RMS differences between the experimentally measured IMU orientations and the virtual IMU orientations. From here we refer to these RMS differences as *inverse kinematics orientation differences*. In the inverse kinematics algorithm, we downweighted the terms corresponding to the distal IMUs (reduced the relative weighting on the tibial IMUs and the foot IMUs) to minimize the influence of the IMUs that were closer to the in-ground metal force-plates.

### Assessing IMU orientation data and joint kinematics

As noted above, the data were collected in a laboratory environment with ferromagnetic disturbances which resulted in distortions in IMU orientation estimates, especially in the heading direction. These erroneous IMU orientation estimates led to exaggerated hip adduction, hip rotation, and ankle flexion in the downstream inverse kinematics solution. To address this, we developed two exclusion criteria to remove low quality IMU orientation data that did not assume knowledge of the true IMU orientations. The exclusion criteria were based on the inverse kinematics orientation differences described above and were as follows: (i) if the differences exceeded a threshold of 45 degrees in the first 10s of the trial, indicating poor tracking of the IMU orientation, then these data were excluded or (ii) if the average range of the difference over 60ms bins (in the first 10s of the trial) exceeded a threshold of 30 degrees, indicating unrealistic variability and therefore poor data, then these data were excluded. We share a subject information table indicating which IMUs were included in our analysis of joint kinematics (Supplementary Table S1). Seven of 11 subjects had at least one IMU excluded from analysis.

### Statistics

We compared the joint angles computed with IMU orientations to the joint angles computed from optical motion capture using root mean square (RMS) difference. We ran a Pearson correlation for each subject and joint angle, and we reported the mean and standard deviation for the correlation coefficients and average change in correlation coefficients over 10 minutes (Table 1). Bilateral joint measures were pooled for all summary statistics. As some of the data were not normally distributed, as determined by a Shapiro-Wilk test, we computed the median and interquartile range of RMS difference over all subjects and all minutes for each joint angle. Outliers were defined as values 1.5 times the interquartile range below or above the 25th and 75th percentile (corresponding to the bottom or top of the box, respectively, in box plots).

We quantified drift for each subject and joint angle using a linear regression on each subject’s individual per-minute RMS differences for each joint angle over the 10 minutes. For each joint angle, we averaged the slopes of the linear fits across subjects. We report the range of slopes to represent drift as an average change in RMS difference per minute. To examine changes over time in RMS differences for different sensor fusion algorithms, we subtracted a subject’s joint angle RMS difference at the end of the first minute from the RMS difference at the end of the 10th minute. We completed all statistical analyses in MATLAB R2019a.

## Results

Median RMS differences between IMU and optical-based kinematics were 3-6 degrees over all subjects and all minutes (Figure 1) for all joint angles except hip rotation (12 degrees); these values are within the reported variability and uncertainty of optical motion capture (35). We saw a similar range of RMS differences between IMU and optical-based kinematics for the 10-minute sequence of lower-extremity movements. Results for the two open-source complementary filters were similar, so we focus on the results produced with the open-source algorithm from Madgwick et al., 2011 (5) and refer to it as the “complementary filter”. Minute-by-minute RMS differences for the complementary filter from Mahony et al., 2008 (3) can be found in Supplementary Figure S2 and highlight that these trends for RMS differences over time were also largely flat.

**Figure 1.**
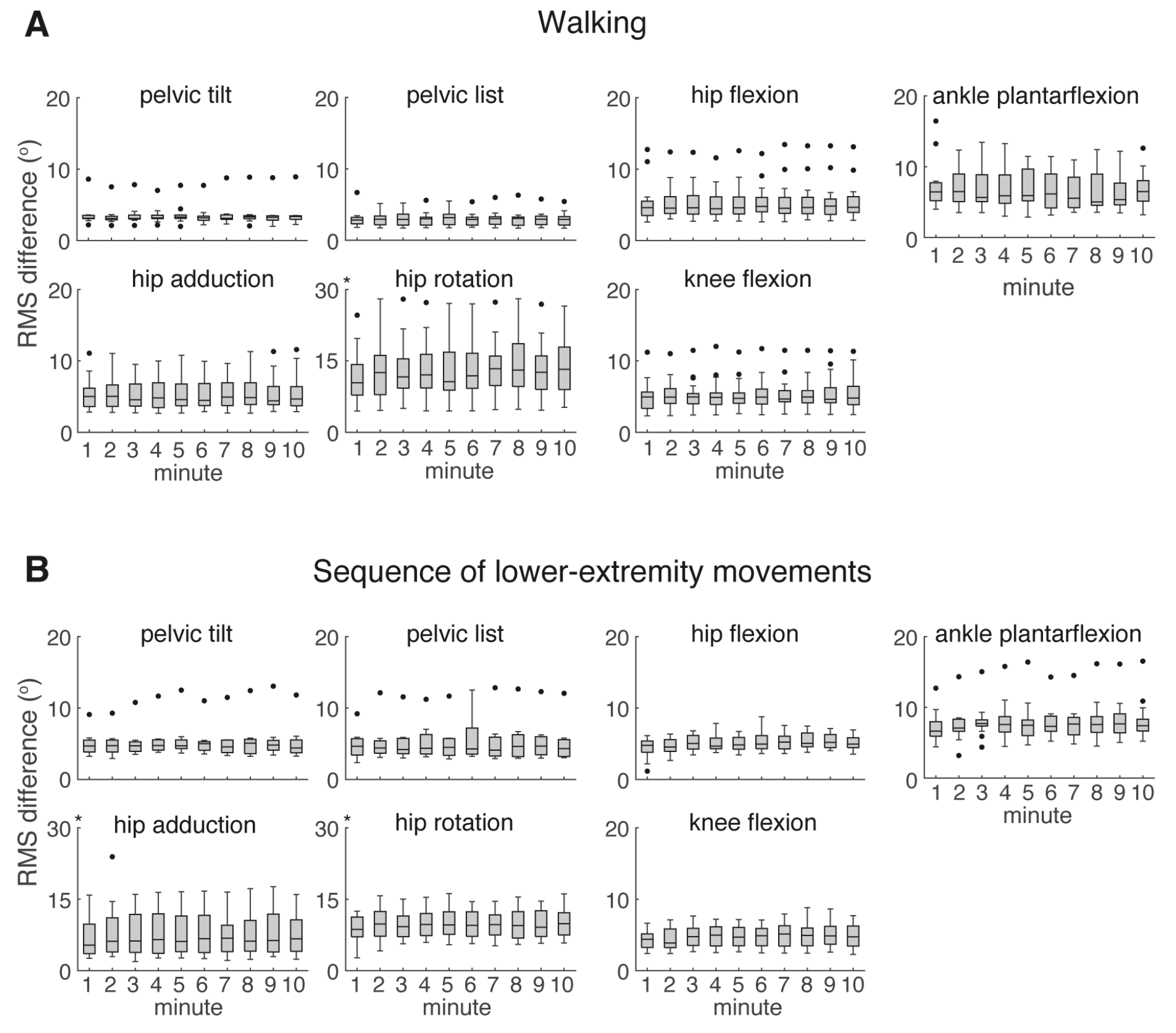
Root mean square (RMS) differences for IMU-based lower extremity joint kinematics over 10 minutes. Our open-source workflow produced IMU-based kinematics comparable to optical-based kinematics during (A) a 10-minute period of overground walking and (B) a 10-minute sequence of common lower extremity movements. Median RMS differences between IMU and optical-based kinematics were 3-6 degrees for all joint angles except hip rotation (12 degrees) over all subjects and all minutes. Flat trends across median per-minute RMS differences highlight minimal drift over 10 minutes. Box plot height is equal to interquartile range with outliers (black dots) defined as values exceeding double the interquartile range. The asterisk * denotes a different y-axis range. Results shown used the complementary filter (5).

Lower-extremity joint kinematics showed minimal drift. Median RMS differences were largely unchanged over 10 minutes for all joint angles during both conditions (Figure 1). Each individual subject’s per-minute RMS differences showed minimal change over 10 minutes for all joint angles (Supplementary Figure S3), and the average slopes of the linear fits to these data were near zero (−0.14 to 0.17 deg/min), indicating minimal drift. Though the linear fits were not strong (R^2^ = −0.03 to 0.4), this was likely due to the nearly horizontal trajectories of the subjects’ RMS differences, at which point R^2^ approaches a negative value (i.e., the chosen model fits worse than a horizontal line).

Individual subjects’ mean IMU-based joint angles over the gait cycle showed minimal difference (within two standard deviations) from optical-based joint angles between the 1st and 10th minute of overground walking (Figure 2). IMU-based hip rotation showed the least agreement with optical-based kinematics. A few subjects (Subjects 2,3,11) had IMU-based kinematics outside two standard deviations of optical-based kinematics (Figure 2).

**Figure 2.**
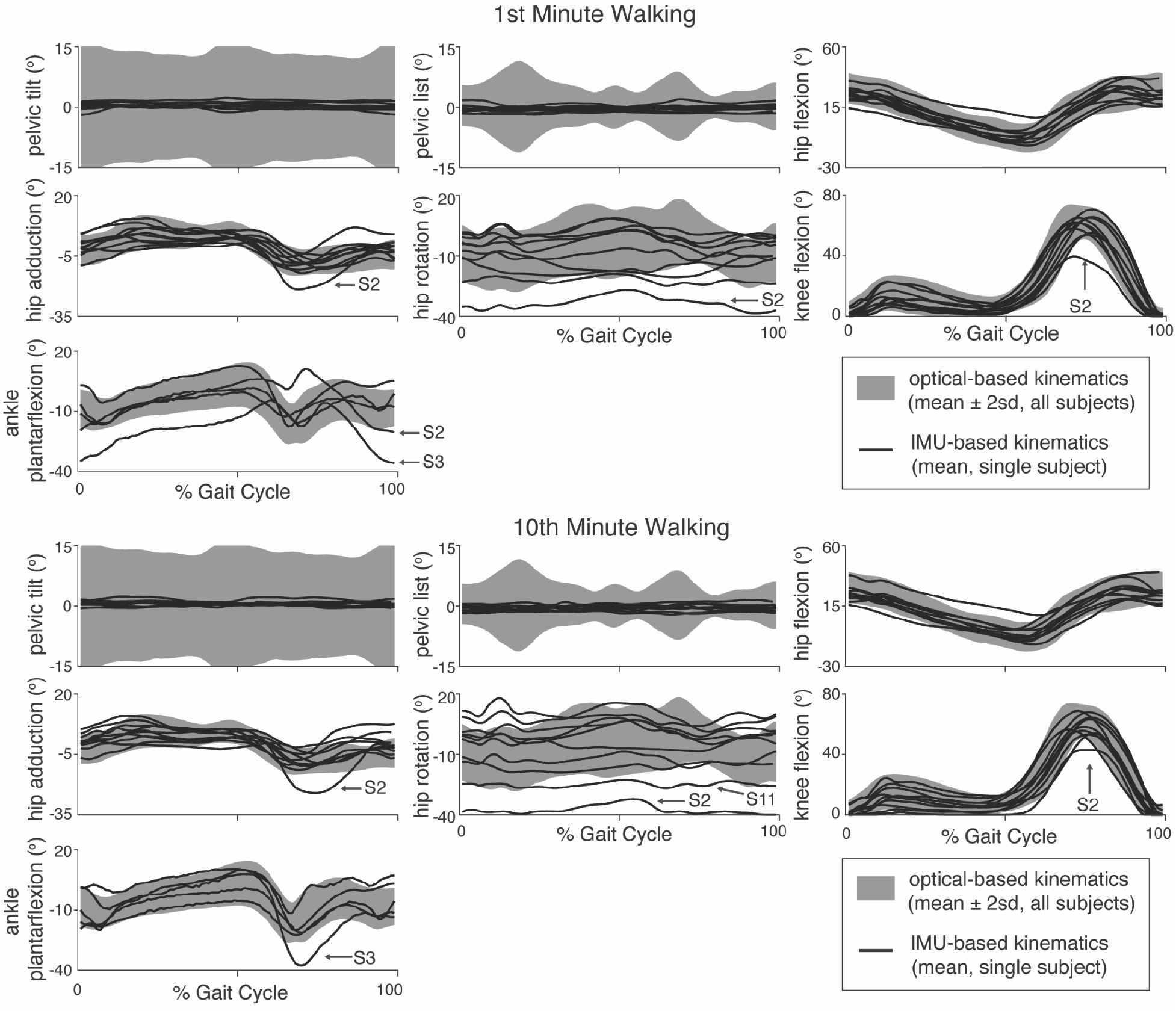
IMU-based lower extremity joint kinematics in the 1st minute (top) and 10th minute (bottom). Individual subjects’ IMU-based kinematics showed minimal drift and agreement with optical-based kinematics between the 1st and 10th minute of overground walking (N = 10 subjects, one subject, S1, lacked any periods of straight-walking and was omitted from this plot). Mean ± two standard deviations (sd) for optical-based kinematics is shown as a grey shaded band and individual subject means for IMU-based kinematics are shown as black lines. Only right-side kinematics are shown, except for one subject (Subject 5) whose right IMU sensors had a technical issue and left body kinematics were used instead. Subjects whose IMU-based kinematics were outside of the optical-based kinematics mean plus or minus 2sd are labeled to demonstrate how kinematics compared for these subjects over time and across joint angles. For example, a magnetometer distortion affected Subject 3’s calibration which affected ankle kinematics in the first minute, but this was resolved by the 10th minute.

We found moderate to strong correlations between the IMU-based kinematics and the optical-based kinematics as indicated by high correlation coefficients over the 10-minute period of overground walking, ranging from r = 0.60 to 0.87 (Table 1). The average difference in correlation coefficient between the first and 10th minute was also near zero (−0.1 to 0.1), indicating little change or drift over the 10 minutes. The results were similar for the sequence of lower-extremity movements (Supplementary Table S2).

**Table 1.** Correlation coefficients between IMU- and optical-based kinematics over 10 minutes of overground walking, averaged over all subjects.

| Joint Angle | Overall Correlation (r) | Average Difference in Correlation (r) Between 1st & 10th Minute |
| --- | --- | --- |
| pelvic tilt | 0.60 (0.30) | 0.1 (0.1) |
| pelvic list | 0.65 (0.23) | 0.002 (0.1) |
| hip flexion | 0.84 (0.29) | -0.002 (0.01) |
| hip adduction | 0.60 (0.27) | -0.002 (0.06) |
| hip rotation | 0.71 (0.27) | -0.03 (0.04) |
| knee flexion | 0.87 (0.32) | -0.003 (0.04) |
| ankle plantarflexion | 0.70 (0.10) | -0.1 (0.1) |

Table entries are: mean (standard deviation).

Joint angles computed using the complementary filter and the proprietary filter from Xsens showed similar changes in median RMS difference over 10 minutes, with less than 2 degrees versus less than 4 degrees change in RMS difference over 10 minutes, respectively, during walking (Figure 3). Similar results were achieved for the sequence of lower-extremity movements (see minute-by-minute RMS differences in Supplementary Figure S4). The proprietary filter, however, had more and larger outliers than the complementary filter (almost 50 degrees change in RMS difference in knee flexion for one subject, Figure 3), highlighting that when the proprietary filter starts to drift, errors can accumulate quickly and substantially.

**Figure 3.**
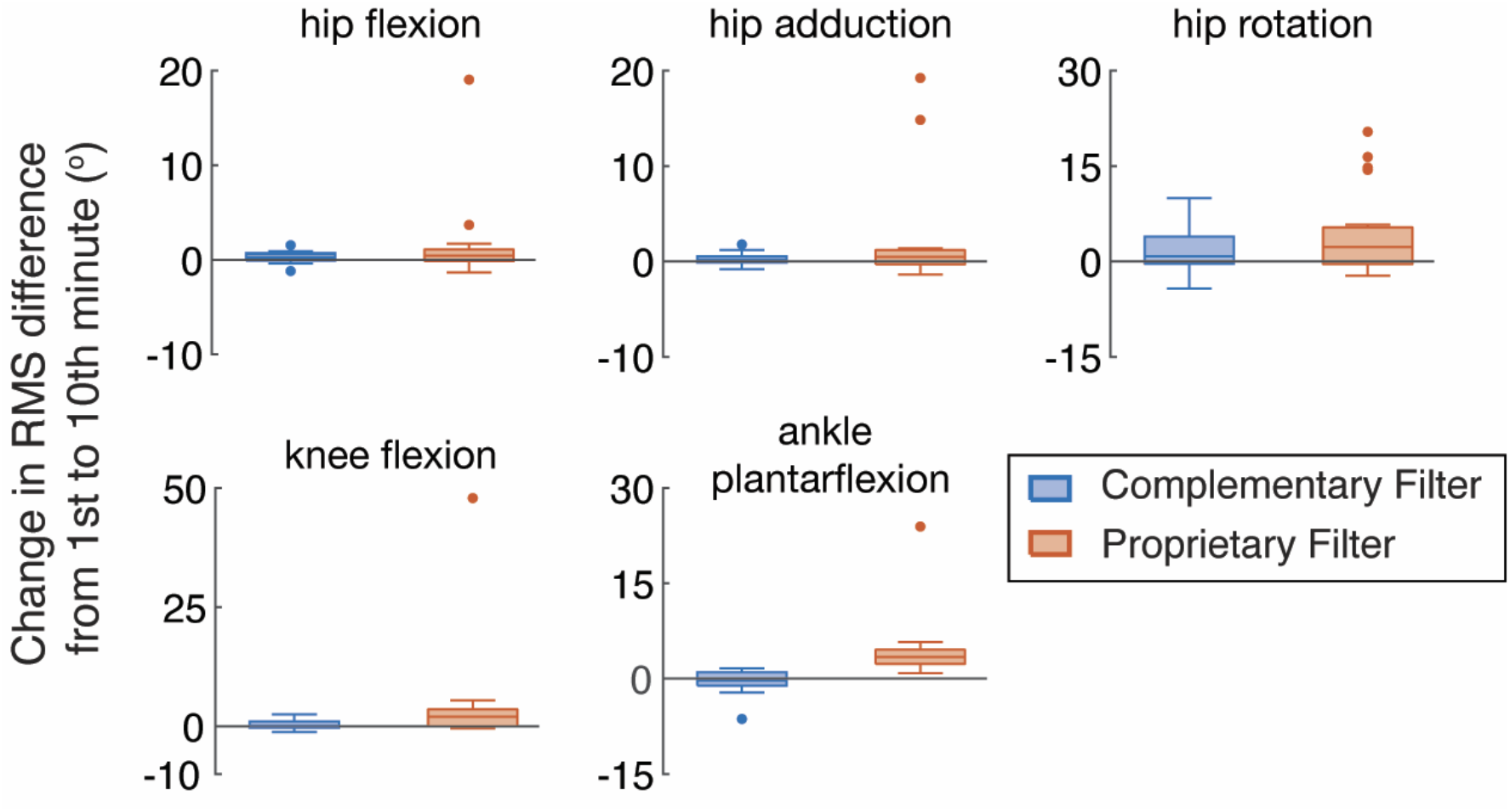
Influence of sensor fusion algorithm on IMU-based lower extremity joint kinematics. The complementary filter (5) outperformed the proprietary filter included with the Xsens sensors in minimizing the change in RMS difference over time, as indicated by the larger changes in root mean square (RMS) differences for the kinematics computed with the proprietary filter compared to those computed with the complementary filter from the first to the 10th minute of overground walking. Outliers from the proprietary filter (orange dots) show that when the proprietary filter starts to drift, errors can accumulate quickly.

We found that down-weighting the distal sensor orientations (reducing the relative weighting on the tibial and feet sensor orientations) when solving inverse kinematics improved the accuracy of the kinematic estimates and reduced drift. The inverse kinematics computed with downweighted distal sensors’ orientations showed less RMS difference (up to 28% less) than inverse kinematics computed with uniformly weighted orientations in the 10th minute of overground walking (Figure 4). Note that all other figures with IMU-based kinematics show downweighted results.

**Figure 4.**
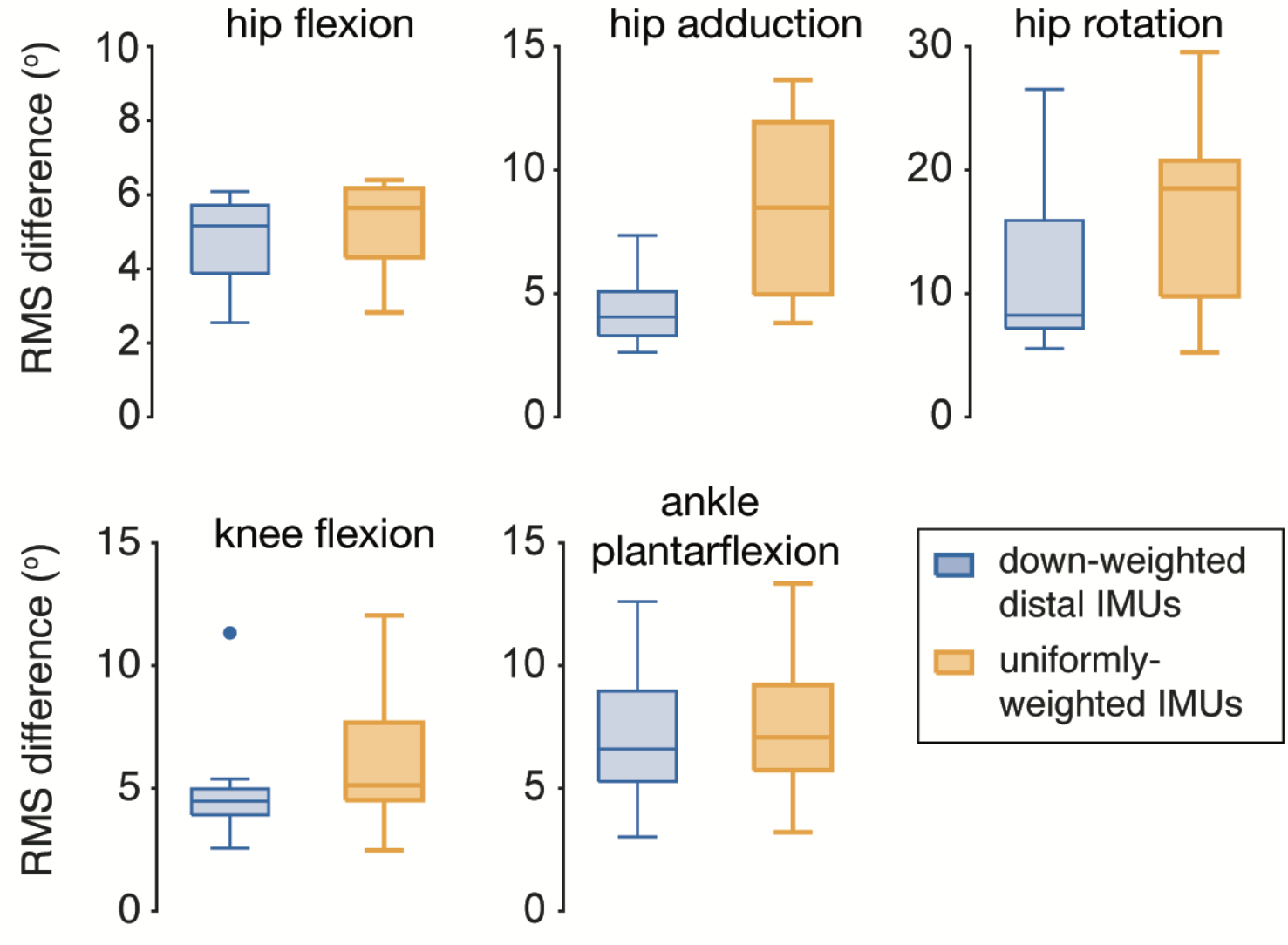
Effect of downweighting distal IMU sensors when solving inverse kinematics. Reducing the relative weighting on the tibial orientations and the calcaneal orientations when solving inverse kinematics helped reduce error as indicated by lower mean joint angle root mean square (RMS) difference in the 10th minute. To highlight how this downweighting influences all joint kinematics, this analysis includes mean joint angle RMS differences for the four subjects who did not have IMUs excluded and results computed from the complementary filter. These trends were also seen with all 11 subjects and with the results computed with the proprietary filter.

Changes in inverse kinematics orientation differences (the angle difference between the experimental IMU orientation and the virtual IMU orientation) from the 1st to 10th minute were strongly correlated with changes in sensor fusion error (Figure 5), indicating that inverse kinematics orientation differences are a helpful tool for tracking errors in the orientations from sensor fusion when present.

**Figure 5.**
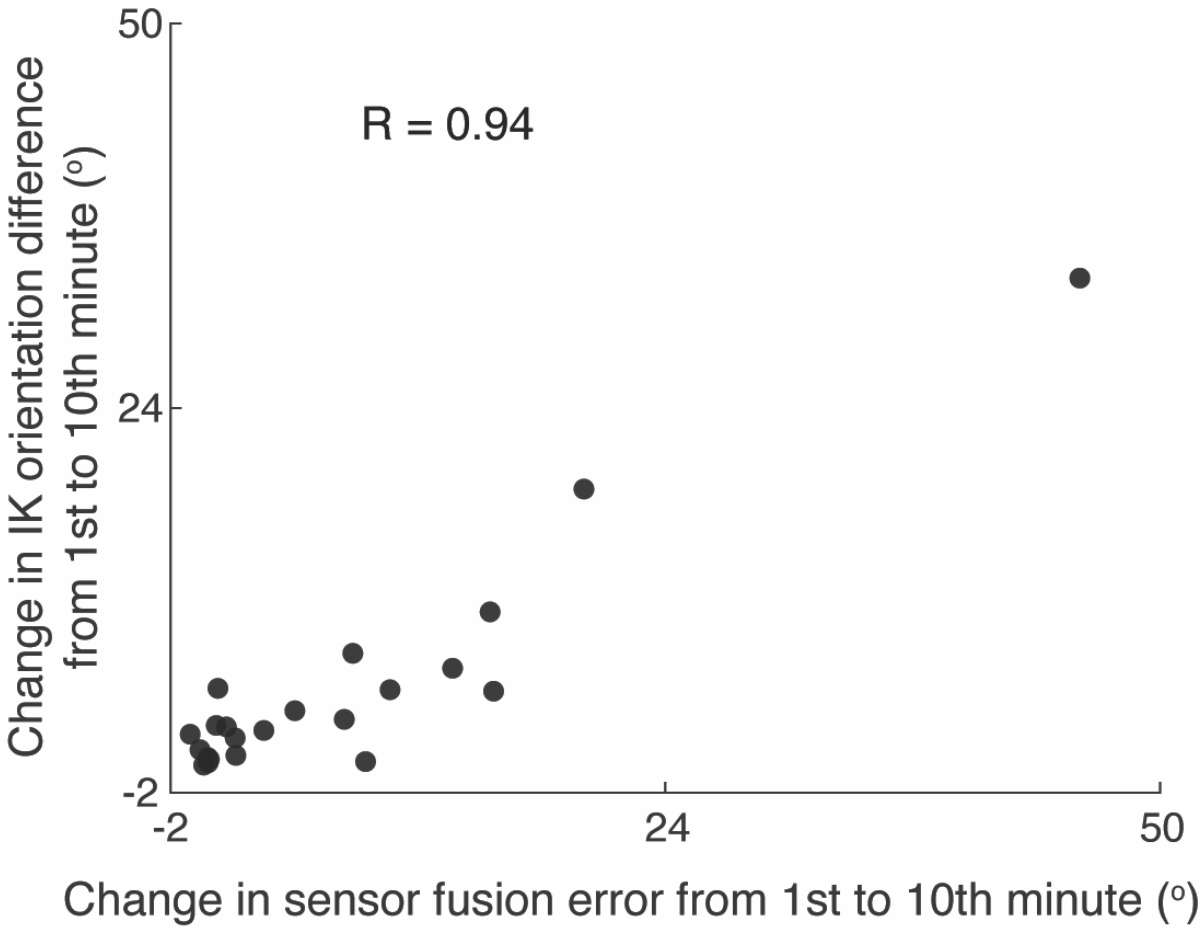
Changes in inverse kinematics (IK) orientation differences relate to changes in sensor fusion errors. Changes in IK orientation differences (mean over all joint angles per subject) from the 1st to 10th minute were strongly correlated with changes in sensor fusion error, indicating that IK orientation differences are a helpful tool for tracking error in the sensor fusion orientation when present. Individual subjects’ data are represented by black circles, and kinematics computed with both the complementary and the proprietary filter were used.

## Discussion

Our open-source workflow for computing three-dimensional lower extremity joint kinematics with IMUs produced joint angles that were consistent with optical motion capture (3-6 degrees RMS differences for all joint angles except hip rotation) and showed minimal drift during a 10-minute period of common lower-extremity movements (including walking). The differences between IMU-based kinematics and optical-based kinematics are similar to previous studies (16), despite our experiments being an order of magnitude longer in duration. We also found that using sensor fusion approaches, as well as downweighting distal IMUs during inverse kinematics, mitigate drift during these common lower-extremity movements. Our open-source workflow, documentation, data, and models are shared at https://simtk.org/projects/opensim so that others can reproduce and extend our work.

Our results suggest that explicit sensor recalibration (e.g., sitting or standing still for a few seconds or zero velocity updates) may not be necessary to mitigate drift when monitoring human movement with IMUs. Our workflow produced results with minimal drift by leveraging complementary filters (3,5) that incorporate magnetometer data. Our study indicates that natural human movement—even continuous walking without explicit periods of rest—includes phases where each individual body segment has a low angular velocity. During these phases, the sensor fusion algorithms of Madgwick and Mahony were able to reject drift. These implicit corrections by the sensor fusion algorithm occurred at different times depending on the activity, but for each of the movement trials (continuous walking and a sequence of movements), the frequency and the duration of periods of low angular velocity were sufficient to mitigate drift. The approach further has the benefit of being activity-agnostic, compared to some previous approaches that were tailored to specific activities (e.g., (36) for running; (37) for specific phases during walking) or reliant on achieving relatively high joint center accelerations (23,38). We also verified that by selecting parameters for the sensor fusion algorithm that mitigated drift, we did not sacrifice accuracy over short durations. In particular, the RMS differences during the first 15 seconds of walking were similar between our sensor fusion estimate and an estimate obtained by solely integrating the angular velocities over time (Table S3). Future work should determine whether the sensor fusion algorithm is sufficiently robust for periods of continuous running, sprinting, or other highly dynamic activities.

The proprietary filter included with the sensors resulted in more drift for all subjects (Figure 3), likely because the filter rejected the distorted magnetometer signal over too long a duration. In fact, it has been shown that the stability of a Kalman filter, on which the proprietary filter is based (24), is time-limited in heavily disturbed magnetic environments; more specifically, accuracies of 3-5 degrees can only be maintained for about 30 seconds in a disturbed magnetic field. However, Kalman filters can recover if there is sufficient time spent in areas with fewer magnetic disturbances though it can take up to 50-60 seconds before orientation estimation error decreases to 5 degrees (18). During the overground walking task, our subjects were walking in a relatively small motion capture volume that included walking over metal, in-ground force plates, and thus little chance for the Kalman filter to recover. While there were some subjects for whom the proprietary filter performed particularly poorly (outliers in Figure 3), it is possible that this could be monitored with real-time inverse kinematics or that results could improve with software/firmware updates from Xsens.

The biomechanical model and inverse kinematics algorithm used in our workflow (OpenSense, OpenSim), provided features that helped us to monitor errors when they did occur. For example, we saw that a change in inverse kinematics orientation differences from the first to the 10th minute was strongly correlated with sensor fusion error (Figure 5) demonstrating utility for monitoring joint angle accuracies. We also used the inverse kinematics orientation differences to exclude IMUs that presented large differences early in the experiment (within the first 10s). Monitoring inverse kinematics orientation differences could alert users to the presence of error, as it does not require knowing the true orientations of the IMUs. This is a salient feature because there has been no standard method to monitor error over time for IMU-based kinematics. As the reliance on IMUs for human movement experiments continues to increase, users will benefit from having error-monitoring features integrated with user-friendly software (OpenSim) that possesses a significant community of users, 25,000+ users/year worldwide (30).

Our study also provides insights about how different joints are affected by magnetic disturbances and drift. For example, while magnetic disturbances can cause large errors in heading angles, our results suggest that these errors do not significantly impact the accuracy of the knee flexion angle. This is an important observation as a major concern in the adoption of IMUs in biomechanics and physical rehabilitation is the presence of magnetic disturbances in the patient’s home. We suspect that knee flexion accuracy is not severely impacted by magnetic disturbances because these disturbances will likely impact the heading of both the shank and the thigh similarly, and possibly because of the kinematic constraint on the knee (i.e., one dof in the sagittal plane). We observed that the IMU orientations computed from sensors on the feet were most affected by magnetic disturbances. In laboratories similar to ours with in-ground force plates, researchers have reported significant magnetic disturbances to the point where they recommended that measurements be performed at least 40 cm off the ground (18). We were able to increase the agreement between IMU- and optical-based kinematics by reducing the frequency at which the complementary filter incorporated magnetometer data for the foot IMU (Supplementary Figure S5). This balanced the positive influence from some magnetometer information and distortion from the force plates. As this may be overfitting to our data, we did not use this approach when reporting our final results, but offer it as an approach to explore further when computing IMU-based kinematics in disturbed magnetic environments.

Similar to past studies, we saw the largest joint angle RMS differences in hip rotation (12.7 degrees median RMS difference). This could be partially due to the fact that the foot IMUs were most affected by ferromagnetic disturbances, and their distorted heading estimates could have introduced error into the hip rotation angle, which was the degree of freedom that relied most on this heading information. RMS errors on the order of 10 degrees are also observed for optical motion capture, which may also have contributed to the RMS differences we observed (35). We also qualitatively observed large hip adduction errors due to magnetic disturbances while the subjects were sitting in the sequence of lower activity movements. Future work is needed to assess joint-specific impacts of magnetic disturbances.

It is important to consider the limitations of our work. We used the optical motion capture data to pose the model for IMU registration, which is an unrealistic approach for natural environments. We instructed subjects to take a “neutral” pose, with pelvic tilt, pelvic list, hip flexion, hip adduction, hip rotation, knee flexion, and ankle flexion at 0 degrees. The mean difference between the subject’s chosen pose (measured with motion capture) and the instructed pose was relatively small (3.8 degrees), and the difference range was 0 - 20.1 degrees (maximum was hip rotation) over all joint angles, and all subjects. A range of calibration approaches including manual, static, functional, and anatomical methods have been described and assessed (7). While using an IMU-based calibration might have altered the RMS differences, we expect our conclusions showing minimal drift would be unchanged.

Despite being one of the longer validation studies for IMU-based kinematics, our 10-minute experiments may not be sufficient to understand the accuracy over multiple hours or days. However, we have three cases where the IMUs were calibrated between seven and twelve minutes before the start of the 10-minute experiment and we again observed minimal drift (Table S4). This suggests that our results might translate to longer durations, but future experiments should assess this. In this study, we focused on assessing the accuracy over an aggregate of activities. Future work should validate the approach for the individual activities in the sequence of lower extremity movements, along with other activities of daily living, including upper body kinematics, and more natural sequences of these activities. Since our sample size was small and our subject demographics were homogenous, it is uncertain how our results may translate to other populations. We hope that future studies will be conducted with larger, more diverse populations and will build upon the data repository we have provided.

## Conclusions

Our open-source workflow (OpenSense, OpenSim) provides accurate estimates of human joint kinematics with wearable technology by leveraging the advantages of inertial sensors, sensor fusion algorithms, and model-driven simulation. The validation over 10-minute durations during common human movements gives confidence to users in being able to monitor and compute kinematics with minimal drift using IMUs. Though all of our kinematics were calculated offline, a recent study has shown promising accuracy over short durations with a low-cost and portable system utilizing the same open-source tools to compute inverse kinematics described here (39). Integration with the OpenSim musculoskeletal simulation environment opens the gateway to investigate other quantities, like muscle mechanics and inverse dynamics with IMUs. Future work will focus on developing algorithms to estimate kinetics from IMU data, and streamlining real-time systems to enable biomechanical monitoring, feedback, and interventions outside of the lab. This suite of open-source tools brings the field closer to conducting human movement studies in natural environments.

## List of abbreviations

(IMU): inertial measurement unit
(IK): inverse kinematics
(RMS): root mean square

