## Supplementary File for "OpenSense: An open-source toolbox for Inertial-Measurement-Unit-based measurement of lower extremity kinematics over long durations"

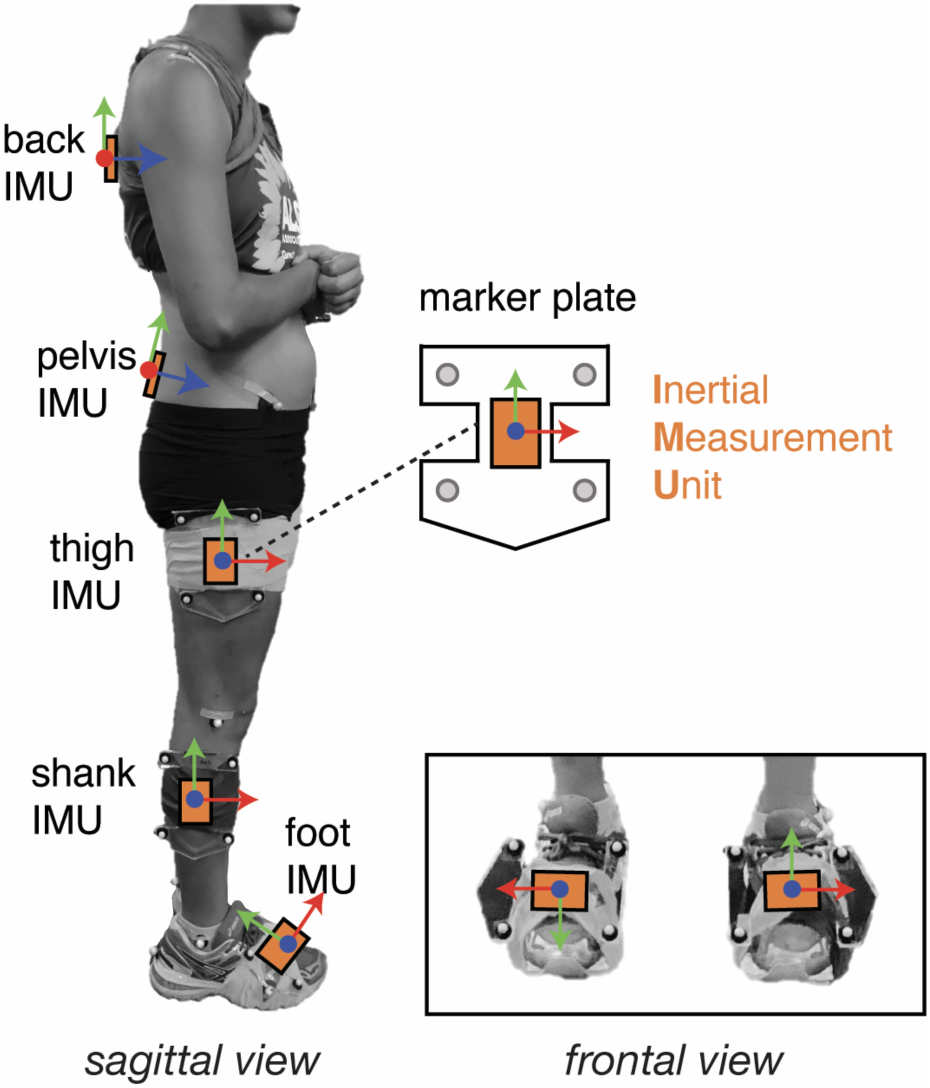


**Figure S1. Experimental setup for marker plate and IMU placement.** Subjects were outfitted with 8 IMUs (MTw Awinda, Xsens North America Inc., Culver City, USA), affixed to a thin plexiglass plate along with a cluster of at least 4 retro-reflective markers placed on the upper back (T2), lower back (L5), and the right and left thighs, shanks, feet.

**Table S1. IMU information per subject**

| Subject | IMU sampling rate (Hz) | Excluded IMUs |
|---|---|---|
| 1 | 100 | both feet |
| 2 | 100 | |
| 3 | 100 | |
| 4 | 100 | |
| 5 | 100 | right body (were not transmitting data), left foot; all data stopped transmitting for sequence of lower extremity movements and was excluded |
| 6 | 40 | right foot |
| 7 | 100 | |
| 8 | 100 | left foot |
| 9 | 100 | both feet |
| 10 | 40 | right foot |
| 11 | 100 | both feet |

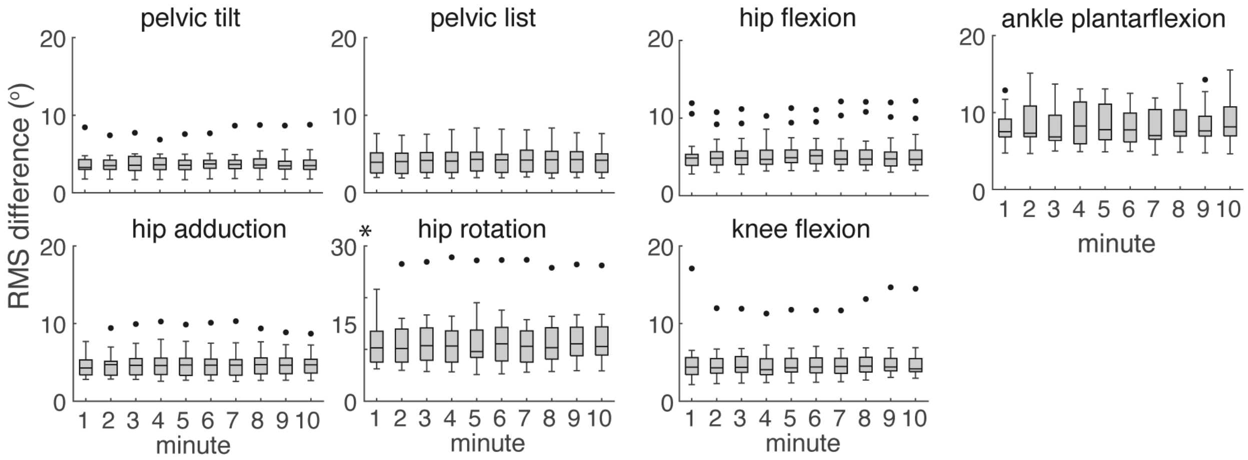


**Figure S2. Errors for IMU-based lower extremity joint kinematics over 10 minutes with the complementary filter from Mahony and colleagues.** IMU-based kinematics computed with a complementary filter (3) over a 10-minute period of overground walking showed minimal drift. Box plot height is equal to interquartile range with outliers defined as data exceeding double interquartile range. The asterisk * denotes a different y-axis range.

**Table S2.** Correlation coefficients between IMU- and optical-based kinematics over 10 minutes during repetitions of a sequence of lower-extremity movements.

| Joint Angle | Overall Correlation (r) | Average Difference in Correlation (r) Between 1st & 10th Minute |
|---|---|---|
| pelvic tilt | 0.51 (0.30) | -0.2 (0.4) |
| pelvic list | 0.75 (0.32) | 0.01 (0.07) |
| hip flexion | 0.98 (0.01) | -0.005 (0.02) |
| hip adduction | 0.75 (0.18) | -0.002 (0.06) |
| hip rotation | 0.63 (0.20) | 0.06 (0.07) |
| knee flexion | 0.99 (0.00) | -0.0007 (0.007) |
| ankle plantarflexion | 0.68 (0.25) | 0.009 (0.02) |

Table entries are: mean (standard deviation).

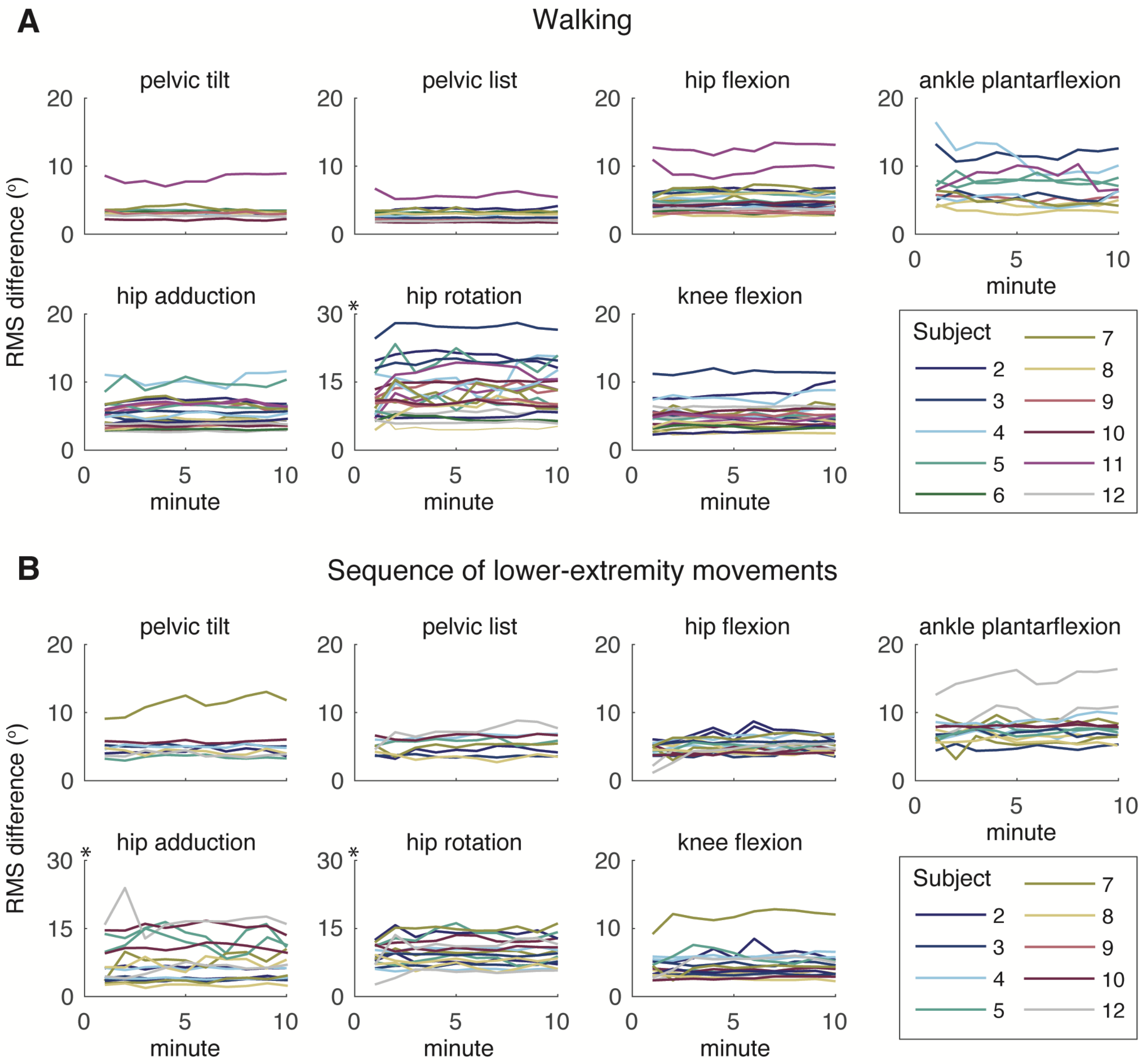


**Figure S3. Minute-by-minute root mean square (RMS) difference trajectories by subject demonstrate minimal change or drift over 10 minutes for (A) a 10-minute period of overground walking and (B) a 10-minute sequence of common lower extremity movements.** Different colored lines represent individual subjects' per-minute RMS differences for unilateral joint angles (e.g. both left and right knee flexion are shown for each subject). The asterisk * denotes a different y-axis range. Results shown used the complementary filter (5).

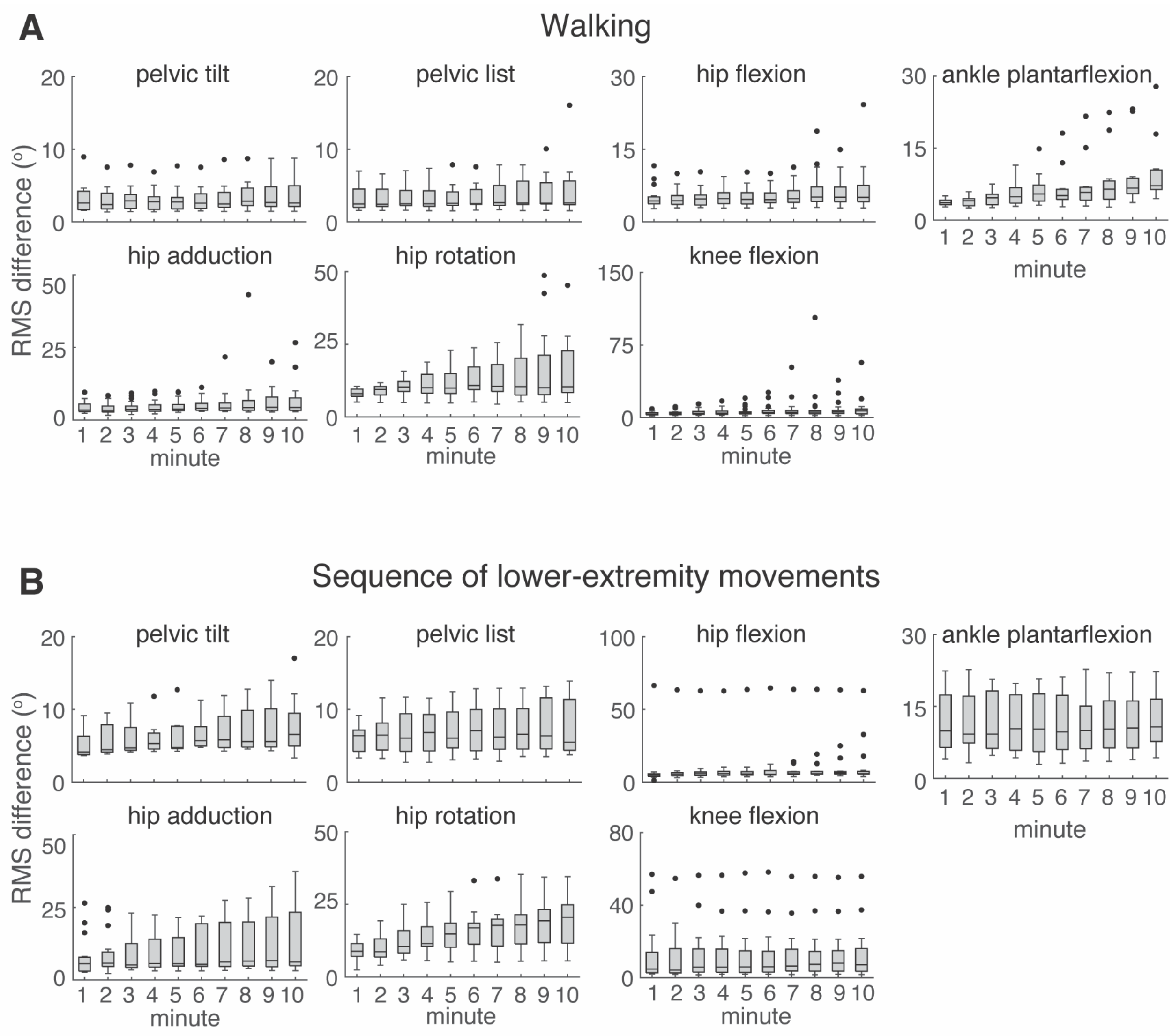


**Figure S4. Errors for IMU-based lower extremity joint kinematics over 10 minutes with a proprietary filter (Xsens).** Root mean square (RMS) differences were less than 4 degrees over (A) a 10-minute period of overground walking and (B) a 10-minute sequence of common lower extremity movements. Box plot height is equal to interquartile range with outliers defined as data exceeding double interquartile range.

**Table S3. RMSE on a short trial of 15s without and with drift correction**

| | Without Correction | With Correction |
|---|---|---|
| pelvic list | 2.9 (0.9) | 2.8 (1.1) |
| pelvic tilt | 2.5 (1.0) | 2.1 (1.0) |
| hip flexion | 4.4 (1.2) | 3.8 (1.5) |
| hip adduction | 4.7 (3.6) | 4.8 (3.4) |
| hip rotation | 7.9 (5.0) | 7.6 (5.4) |
| knee flexion | 3.2 (2.0) | 3.7 (2.4) |
| ankle plantarflexion | 5.3 (3.2) | 6.6 (6.1) |

Table entries are: mean (standard deviation).

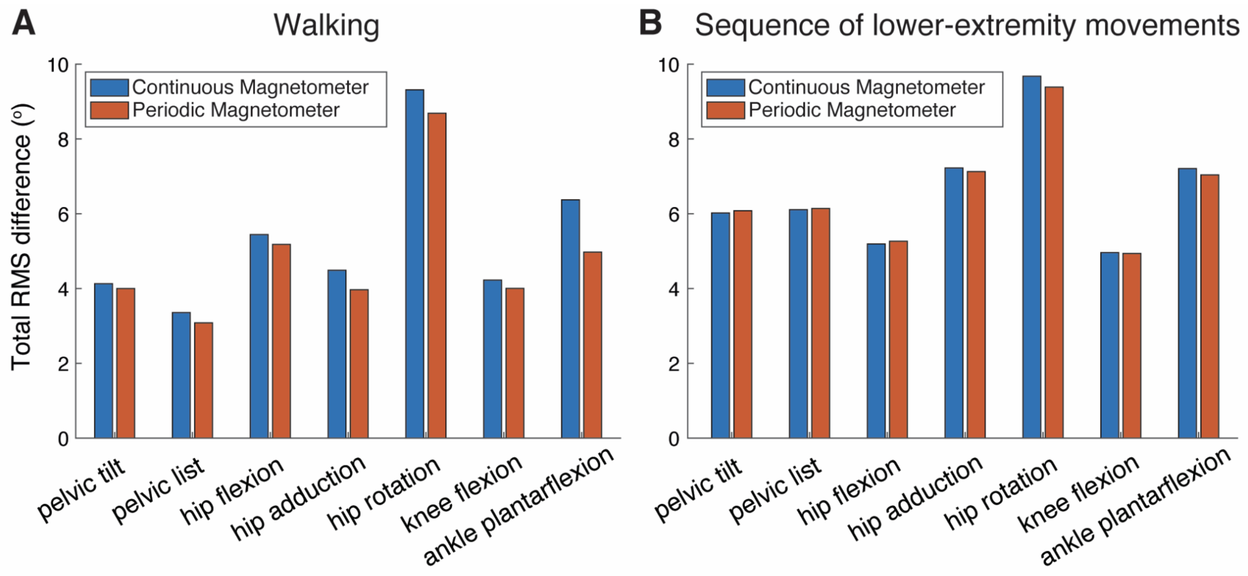


**Figure S5. Periodically weighting the magnetometer reduces errors in joint kinematics.** Periodically, rather than continuously weighting the magnetometer on the foot IMUs reduced the joint total root mean square (RMS) difference especially for ankle flexion. Average RMS differences are shown for all subjects that had at least one foot sensor, (A) six subjects in total during the walking task and (B) four subjects in total during the sequence of common lower extremity movements.

**Table S4. RMS with and without seven and twelve minutes of IMU data running before the start of the 10-minute walking experiment averaged over three subjects.**

| | With Extra Duration | Without Extra Duration |
|---|---|---|
| pelvic list | 4.8 (0.5) | 4.8 (0.4) |
| pelvic tilt | 5.6 (0.8) | 5.6 (0.9) |
| hip flexion | 5.8 (1.2) | 5.8 (1.2) |
| hip adduction | 5.6 (1.4) | 5.7 (1.5) |
| hip rotation | 9.1 (2.1) | 9.1 (1.9) |
| knee flexion | 4.6 (1.3) | 4.6 (1.2) |
| ankle plantarflexion | 6.3 (1.5) | 6.8 (1.7) |

Table entries are: mean (standard deviation).
